# Two views of the brain are reconciled by a unifying principle of maximal information processing

**DOI:** 10.1101/2025.11.25.690580

**Authors:** Leandro J. Fosque, Woodrow L. Shew, ShiNung Ching, Keith B. Hengen

## Abstract

There is selective pressure on brains to maximize computational capacity and adaptability in an unpredictable world. Prior work suggests that this demand is satisfied by a regime called criticality, which has emerged as a powerful, unifying framework for understanding how computation can arise in biological systems. However, this framework has been confined to high-dimensional network models. At first glance, this appears irreconcilable with many of the foundational, low dimensional dynamical models that have driven progress in theoretical and computational neuroscience for a century. If criticality is a universal principle, then all models that accurately capture significant aspects of brain function should be constrained by the same fact. Lacking a definition of criticality in low-dimensional dynamical systems, this has been impossible to evaluate. Here, we develop a mathematical definition of criticality that transcends dimensionality by recognizing temporal scale invariance as analogous to spatial scale invariance that defines criticality in large systems. We demonstrate that there are two mechanistically distinct sources of criticality at bifurcations, one deterministic and one that emerges from noise fluctuations. Further, we show that some but not all canonical bifurcations in neural models exhibit criticality, and only a subset of these are biologically plausible. We conduct numerical analyses demonstrating that information processing capacity peaks at critical bifurcations, and evaluate which historically influential neural models contain these bifurcations. Our results establish criticality as a universal neurobiological principle that is accessible to systems of any dimensionality. This unifies disparate modeling approaches under a single computational framework and suggests that optimal information processing emerges not from model-specific mechanisms but from fundamental properties of critical dynamics themselves.

## Introduction

Brains face a fundamental challenge — they must learn and adapt in an uncertain world. Unlike engineered systems designed for specific tasks, biological neural networks cannot know in advance what computations they will need to perform. This demands a remarkable starting point: some state that maximizes the capacity for arbitrary computation across many spatiotemporal scales. Intuitively, such a starting point must comprise conditions in which small changes may drive large and varied responses. However, the ability to produce a full range of responses is non trivial. Theory of both biological^1–3^ and synthetic computing systems^4,5^ suggests that such versatile, multiscale responses that maximize computational power are often found at the boundary between distinct dynamical regimes for instance, the border delineating stability and instability^6,7^, asynchronous and synchronous dynamics^8–10^, or even two different stable states^11,12^. Most of these examples come from high-dimensional systems (e.g., a large number of neurons) and the boundary between these regimes is called *criticality*. In low-dimensional dynamical systems of single neuron spiking^13,14^ and neural mass models^15,16^, similar boundaries are called *bifurcations*, and include transitions between quiescence and spiking, regular and chaotic firing, oscillations, or distinct bursting patterns. Do versatile, multiscale responses emerge near bifurcations in low dimensional systems, as they do in high dimensional systems near criticality? The similarities are qualitatively obvious, but a unified framework for describing bifurcations and criticality has been lacking. This comprises a powerful disconnect in neuroscience: there are two ways to think about the transitions that may explain the power of the brain. These ideas have not been reconciled due to non-overlapping mathematical and conceptual assumptions. Here we show that temporal properties of low-dimensional systems are analogous to spatial (and/or spatiotemporal) properties of high-dimensional systems, thus reconciling these perspectives and unifying a wide range of influential models with a single principle of maximal information processing.

There are prominent differences between the two prevailing notions of state boundaries in complex systems: phase transitions and bifurcations. Phase transitions are easily illustrated by considering water. At atmospheric pressure, the difference between water at 5° C and 50° C is unremarkable, but a tiny nudge from 99.9° C to 100.1° C produces a dramatic change whereby liquid becomes vapor; this is a phase transition. A subset of phase transitions are particularly remarkable. At 374° C and 218 atmospheres, the spatial structure of water becomes complex and multi-scale with some patches of high density liquid and some patches of steam (Fig. 1A). Here, local molecular interactions cascade across all scales, producing emergent phenomena that cannot be predicted from microscopic rules alone. This is *criticality*, where the system exhibits its most complex behavior. Theoretical models of neural systems that are high-dimensional (i.e., a large number of neurons) often have critical phase transitions^9,17–19^. Such high-dimensional models are responsible for many advances in how we understand the brain. Integrate-and-fire models^20^ revealed how spike timing precision emerges from noisy inputs, forecasting the discovery of temporal coding in sensory systems. The Kuramoto model^21^ described the synchronization of weakly coupled oscillators, explaining phenomena including gamma oscillations^22^ and slow wave activity^23^, later validated empirically. Experiments suggest that the complex, multi-scale structure and dynamics of neural activity emerges because the brain is near criticality^3^. Unlike water, criticality in the brain is homeostatically maintained^24^ and accounts for computational depth by offering efficient coding, maximal dynamic range, information capacity, and sensitivity to inputs^1,12,25,26^.

**Figure 1:**
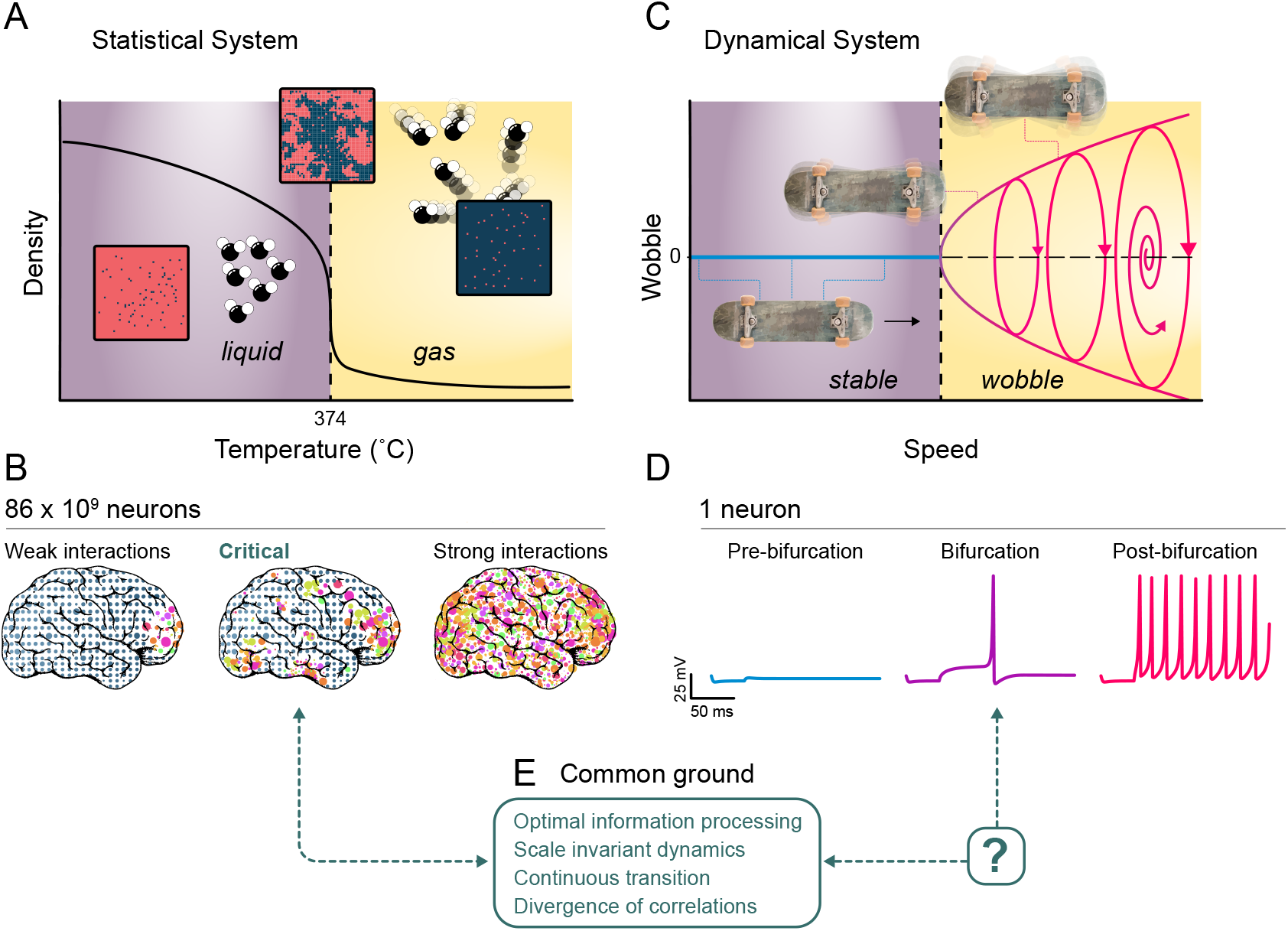
Common ground principles of criticality and bifurcations? **(A)** Phase diagram of water at a pressure of 218 atmospheres. There is a critical phase transition from liquid to gas as the temperature is increased. At criticality (374 C), complex, scale-invariant spatial patterns of low and high density emerge (insets represent spatial patterns of low (blue) and high (red) density). **(B)** Conceptual illustration of a phase transition in an entire brain, where gray dots represent damped and unexcited neuronal populations due to weak interactions between neurons. Conversely, colored dots depict excited populations arising from strong interactions. At the transition between damped and excited, mixtures of gray and colored groups appear at all scales. **(C)** Bifurcation diagram for “death wobbles” on skateboard^27^, where oscillations emerge when speed exceeds a certain value. Below this speed, the skateboard’s motion is stable. **(D)** Conceptual demonstration of single neuron membrane potential considered as a low-dimensional dynamical system. Spikes emerge when current injection crosses a bifurcation. **(E)** Here we seek common ground properties of criticality and bifurcations, focusing on scale invariance and maximized information processing capacity.

The basic definition of a phase transition necessitates a large number *N* of interacting elements (the thermodynamic limit, *N*→ ∞); *N* is the dimensionality and might represent the number neurons in the brain or water molecules in a pot of water. In contrast, bifurcations can occur in a system with only one component (i.e., a single variable). An example of this is the sudden emergence of uncontrollable, side-to-side oscillations of a skateboard at a high enough speed — colloquially known as the death wobbles^27^. Bifurcations are fundamental properties of dynamical systems models that have been the cornerstone of theoretical neuroscience since its inception, often predicting phenomena years before experimental confirmation. Hodgkin and Huxley’s equations^13^ not only explained action potential propagation but predicted the existence of voltage-gated ion channels discovered decades later. Likewise, Wilson and Cowan’s neural mass model anticipated that excitatory-inhibitory balance generates cortical oscillations^15^.

In short, the corpus of theoretical neuroscience leaves us with a zoo of lowand high-dimensional models, each successful in describing a specific aspect of how the brain works, but each tailored for a specific set of applications. Given their collective ability to explain so much of the brain, it is reasonable to ask if effective models, regardless of their dimensionality, are tied together by a common mathematical rule. If so, such a principle would likely be central to a meaningful understanding of the brain. Interestingly, while criticality is defined in high dimensional systems, its central feature — the presence of information across all scales — does not require high dimensionality. In low dimensional systems, this requires only substituting space with time.

Here, we offer four primary contributions. First, we develop a rigorous mathematical framework for criticality in low-dimensional dynamical systems. Second, we develop theory and perform simulations demonstrating that information processing is maximized in dynamical systems tuned to a critical bifurcation. Third, we reveal that only a subset of low-dimensional systems can be critical and biologically plausible. Finally, we examine many historically important models of the brain and conclude that, beyond the level of spike generation, all are unified by possessing a bifurcation that generates criticality. This supports a view that the brain’s computational capabilities may arise not from a collection of disparate mechanisms, but from a universal organizing principle that manifests across all scales of neural organization.

## Results

The reconciliation of low-dimensional bifurcations and high-dimensional phase transitions is achieved by considering time and space on similar footing. Traditionally, an understanding of phase transitions hinges on spatial correlations and spatial patterns, i.e., how neurons in different locations are coordinated. Here we will consider temporal patterns and correlations in low-dimensional systems, and furnish a definition of criticality that does not require spatially extended, high-dimensional systems. Our approach complements recent developments of temporal renormalization group theory, in which temporal scale-invariance defines criticality^28^.

Two key ingredients define criticality^3,29,30^. The first is *scale invariance*; no characteristic or preferred scale of activity dominates. At criticality, activity measurements are statistically similar at different scales and follow a power law (linear on log-log scale). However, scale invariance is insufficient; some instances of scale invariance are unrelated to criticality. For example, word frequencies in a written text tend to be power law distributed (Zipf’s Law). Thus, a second defining criterion is required; this is the notion of a boundary between qualitatively different regimes discussed above. Tuning some control parameter of the system (temperature in the water example, speed in the skateboard example; Fig. 1) across a boundary causes a dramatic, qualitative shift from one regime to another. If a system is 1) at such a boundary and 2) exhibits scale-invariance, then it is at criticality. This comprises a general definition of criticality (formalized in Box 1) which allows us to ask whether dynamical systems — even those representing only a single neuron — can satisfy the definition (Fig. 1E).

Summarily, and as we will show in detail below, this definition identifies two sources of criticality that are mechanistically distinct from traditional phase transitions^31–33^. First, in low-dimensional systems *without* noise, criticality is a consequence of the nonlinearity of system dynamics precisely at the bifurcation. Second, in more realistic low-dimensional systems, i.e., those subjected to some degree of noise, a Brownian (randomwalk) criticality arises at the bifurcation. In biologically plausible systems, such as a brain, it appears inevitable that these two forms co-exist. Below, we identify which bifurcations satisfy this definition of criticality both with and without noise, we ask which bifurcations are biologically plausible, and we analyze their information processing properties.

### Definition 1.

Criticality for Dynamical Systems

Let *ẋ* = *f* (*x, µ*) be a smooth dynamical system with state variable *x* ∈ R^*N*^, control parameter *µ* ∈ R, and fixed points *x*\*(*µ*) satisfying

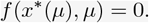

Define the *order parameter* along a fixed point as the function

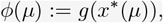

where *g*: R^*N*^→R^*M*^, with *M* ≪ *N* is a smooth observable mapping the microscopic state to a macroscopic quantity. The order parameter captures the emergent collective behavior of the system, such as the population-averaged firing rate or a symmetry-invariant projection of the state *x*.

Suppose that at *µ* = *µ*_*b*_, the system experiences a bifurcation, i.e., a boundary between multiple qualitatively different regimes. Define an open interval Γ ∋ *µ*_*b*_. A dynamical system is **critical** at *µ*_*b*_ with respect to the order parameter *ϕ*(*µ*) if the following condition holds:

**Scale invariance**

There exists a functional *L*[*ϕ*] such that, at the bifurcation *µ*_*b*_, its dependence on an observation variable *O* follows a power law

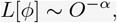

where *α >* 0 is a critical exponent. *L*[*ϕ*] may represent, for example, the relaxation of *ϕ* after a perturbation or the distribution of fluctuations of *ϕ* (e.g., event sizes or durations). When *ϕ* is vector-valued, different components or projections of *L*[*ϕ*] may yield distinct critical exponents. In all cases, the absence of a characteristic scale is captured by asymptotic power-law behavior.

### Which bifurcations are critical?

We consider a broad class of low-dimensional models defined by

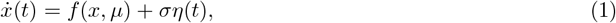

where *f* (*x, µ*) is the deterministic drift, 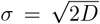 is the standard deviation of the noise with *D* as the diffusion constant, and *η*(*t*) is Gaussian white noise with ⟨*η*(*t*)*η*(*t*′)⟩ = *δ*(*t* − *t*′) and ⟨*η*⟩ = 0. Any model that obeys Eq. 1 — with noise (*σ* ≠ 0) or without (*σ* = 0) — meets the boundary criterion of our definition; a bifurcation, if one exists, emerges only when the control parameter *µ* is tuned to a specific value (*µ*_*b*_ = 0 for the normal form bifurcations considered here). The second defining criterion of criticality — scale invariance — determines whether a bifurcation is critical.

In deterministic systems without noise (*σ* = 0), scale invariance is assessed by considering the response to a stimulus. Specifically, activity *x* is measured in time following a brief perturbation *x*→ *x*\*+ *δx* near a fixed point *x*\*, akin to measuring a peri-stimulus response time series in an experiment (illustrated by *perturbation and result* in Fig. 2 and the time series in Fig. 3). In this context, scale invariance entails a response that decreases according to a power-law *δx*(*t*) ~ *t*^*−α*^(see Appendix A).

**Figure 2:**
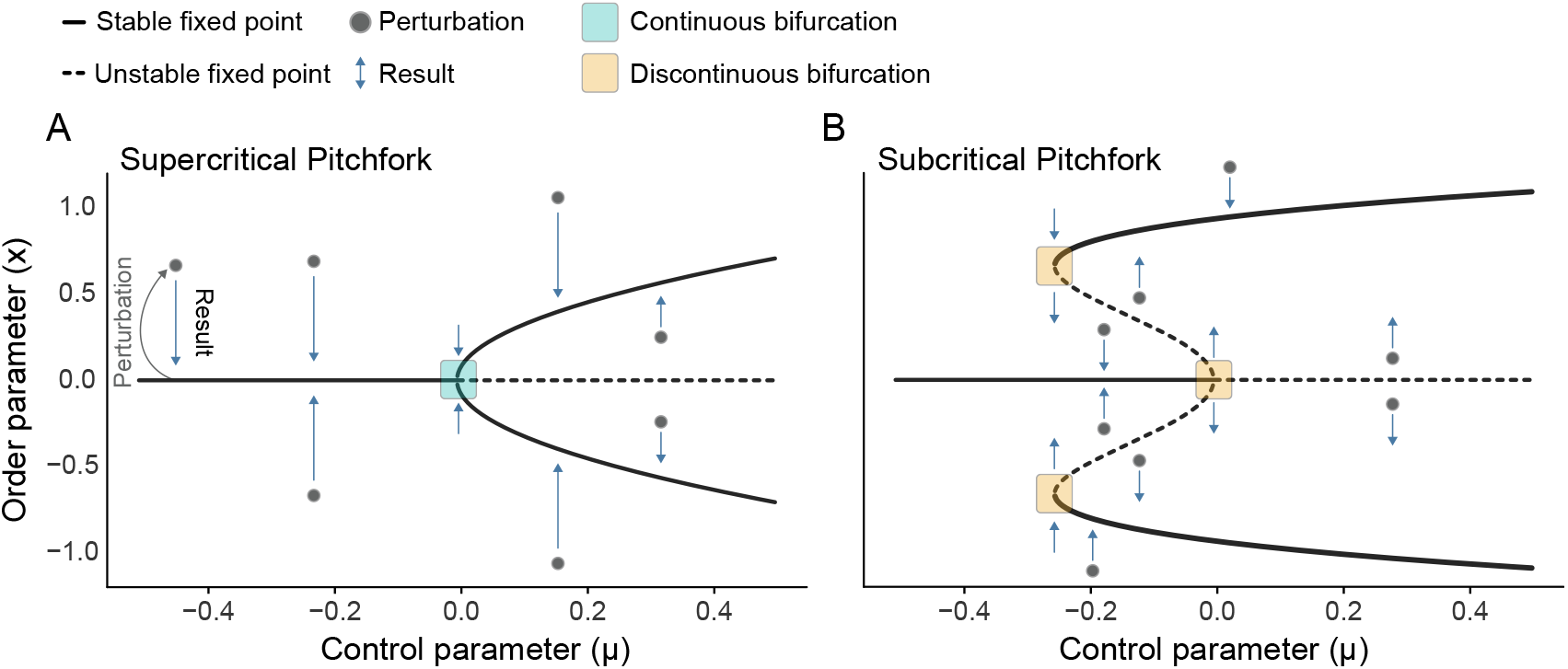
Pitchfork bifurcations. The normal forms of two common bifurcations in dynamical systems are shown with cartoon illustrations to describe their behavior. Fixed points are those locations in the diagram where the system can remain indefinitely; these are illustrated by lines. Solid lines comprise stable fixed points, i.e., fixed points to which the system will return if perturbed. Dashed lines represent unstable fixed points. Here, any perturbation will cause the system to fly away from the fixed point. **(A)** Supercritical pitchfork bifurcation diagram with a series of perturbations (gray dots) and resultant trajectories (blue arrows) to illustrate the behavior of the system around its fixed points. Teal square indicates a continuous bifurcation, where multiple stable fixed points meet. **(B)** Subcritical pitchfork bifurcation diagram with saddle nodes. In contrast to A, all bifurcations are discontinuous (orange square), such that distinct stable fixed points are not in direct contact with one another. The system must jump to traverse these states.

**Figure 3:**
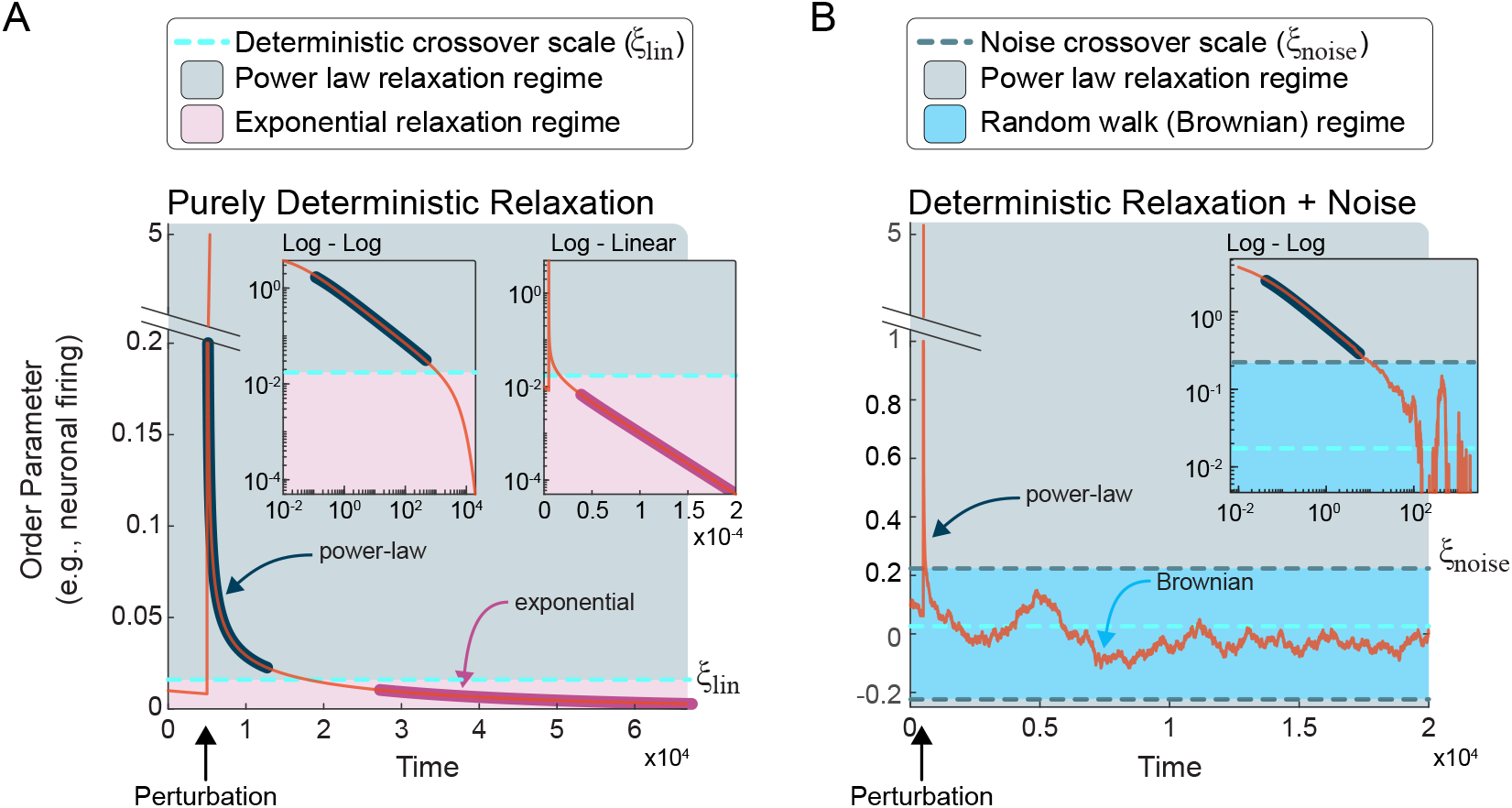
Cross-over scales and two sources of scale invariance. **(A)** Near the bifurcation in a purely deterministic system, perturbations above the crossover scale (*ξ*_*lin*_) relax according to a power law until *ξ*_*lin*_, at which point relaxation becomes exponential. The **left** inset plots the relaxation on log-log axes, in which linearity indicates a power law. Note that the region of the relaxation above *ξ*_*lin*_ is linear (highlighted in navy). The same perturbation and relaxation are plotted in log-linear axes in the **right** inset. Here, linearity is a signature of exponential relaxation, which occurs below *ξ*_*lin*_, highlighted in magenta. **(B)** Additive Gaussian noise results in a second source of scale invariance. Near the bifurcation, perturbations or noise whose amplitude is less than the noise crossover scale (*ξ_noise_*) approximates Brownian criticality, in which the order parameter exhibits a scale-invariant random walk. Above *ξ*_*noise*_, the order parameter relaxes according to a power law, as demonstrated by linearity in the **inset**, highlighted in navy. Note: for each panel, simulations were run on the supercritical pitchfork normal form with *µ* = −3 *×* 10^−4^ and the perturbation comprised kicking the order parameter to *δx* = 5.

Which deterministic dynamical systems exhibit scale-invariant responses to stimuli? The form of *f* (*x, µ*) at the bifurcation is the deciding factor. As detailed in Appendix A, if *f* (*x, µ*_*b*_) can be described in terms of its leading nonlinear term *f* (*x, µ*_*b*_) ≈ *cx*^*n*^, then it will produce a response with power-law decay if either of the following conditions is satisfied:

- *n* is odd and *c <* 0. In this case, the power-law decay is symmetrical (for example, see the bifurcation in Fig. 2A).
- *n* is even and the perturbation arises from the side of the stable fixed points. In this case, the power-law decay is asymmetric; power law decay is only observed when perturbed from one side (for example, saddle nodes, such as the upper and lower bifurcations in Fig. 2B).

These conditions are met by many prevalent bifurcations — pitchfork, transcritical, saddle node, Hopf, and more — and exclude the linear case *f* (*x, µ*) = *µx* as summarized in Fig. 4 and derived in Appendix A. (*n* = 3 for pitchfork and *n* = 2 for transcritical/saddle-node.) For these cases, at the bifurcation, stimulus response is

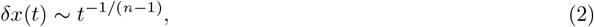

for *n*≥ 2. More generally, when not at the bifurcation (*µ ≠ µ*_*b*_), this scale-invariant response is also present if the nonlinear terms in *f* (*x, µ*) dominate the linear terms, which occurs if the perturbation *δx* is sufficiently large compared to *µ*. As *δx* decreases towards the fixed point there is a crossover, *ξ*_*lin*_, from power-law (for *δx* ≫ *ξ*_*lin*_) to exponentially (for *δx* ≪ *ξ*_*lin*_) decreasing response (Fig. 3A, Appendix A). The precise value of the crossover *ξ*_*lin*_ depends on the specific form of *f* (*x, µ*), but generally *ξ*_*lin*_ approaches zero as *µ* approaches the bifurcation *µ*_*b*_. When *δx* is well below this crossover, the deterministic drift term becomes approximately linear *f* (*x, µ*_*b*_) ≈ *g*(*µ*)*x*. Thus, the time scale of exponential decrease is determined by *µ*. As *µ* approaches *µ*_*b*_, this time scale diverges. This phenomenon is known as *critical slowing down*, and has been of great utility in clinical contexts, as such slowing presages, for example, an impending seizure^34,35^ or sleep transitions^36^. For a purely linear system *f* (*x, µ*) = *µx*, there is no crossover; the response is exponential decrease no matter how large the stimulus. As a result, critical slowing down is insufficient to indicate criticality; critical slowing down occurs in deterministic linear systems, but scale invariance does not.

**Figure 4:**
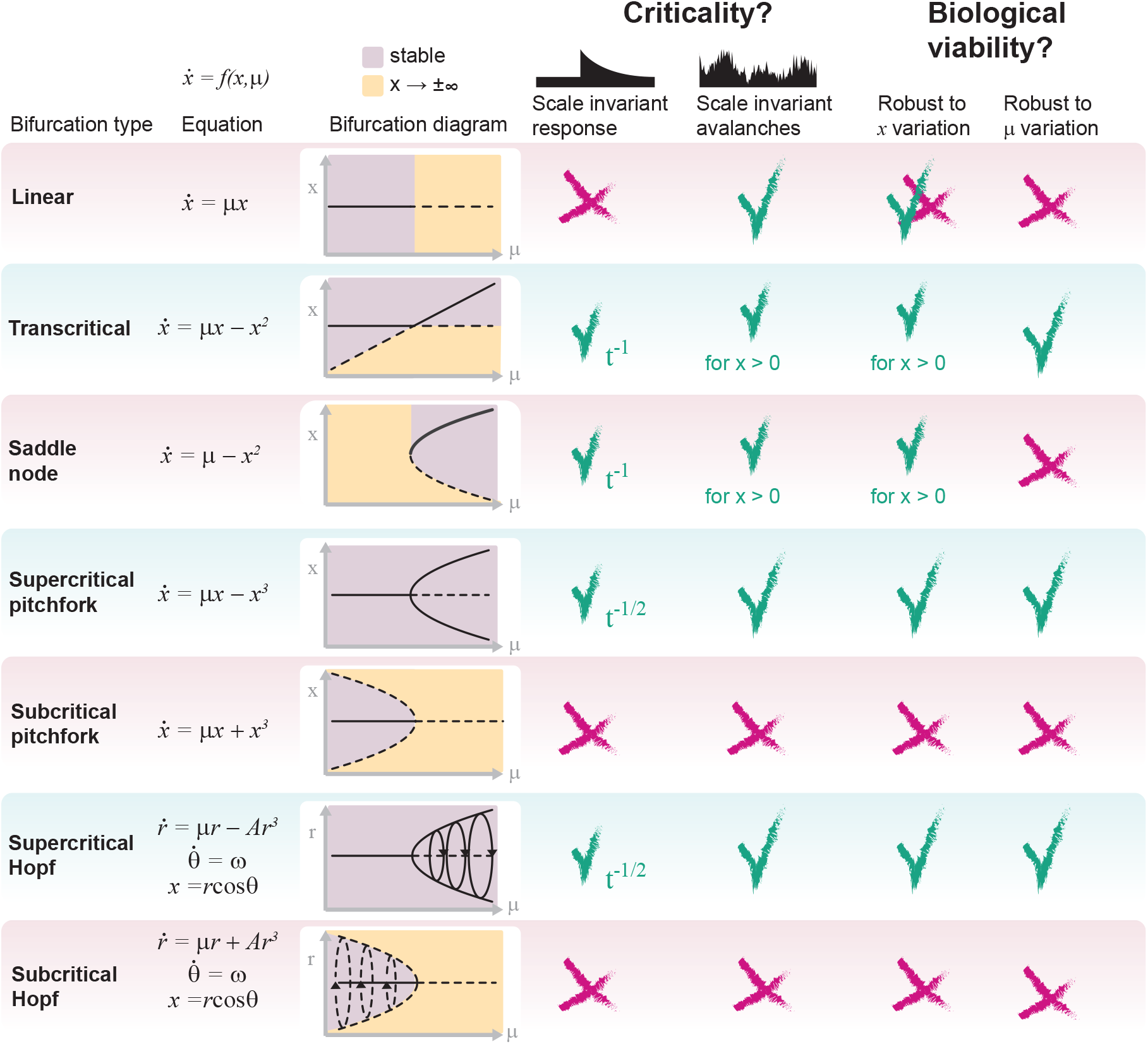
Which canonical bifurcations are critical and biologically plausible? Summary of analyses of seven key bifurcation types. Bifurcation type and normal form (equation) are displayed in the left two columns. The bifurcation diagram is displayed in the third column, and the regions of the diagram corresponding to stable and fly-away (unstable) regimes are shown in purple and gold, respectively. To assess criticality, two sources of scale invariance are analyzed. First, the ability of each bifurcation to support scale-free (power law) relaxations upon a perturbation to the order parameter (*x*, y axis variable) when the control parameter (*µ*, x-axis variable) is tuned to the bifurcation (column 4). Second, the ability to support scale-invariant Brownian (random walk) criticality when subjected to additive Gaussian noise (5^th^ column). Finally, the robustness of each bifurcation to noise/perturbations in the order (6^th^ column) and control (7^th^ column) are evaluated as necessary conditions for biological plausibility.

While mathematically tidy, the noise-free deterministic scenario falls short of describing biological systems, which invariably entail some degree of noise. In other words, in addition to brief perturbations, there are continuous fluctuations in the order parameter; *σ* ≠ 0 in Eq. 1. When accounting for such fluctuations, a second type of scale-invariance arises that is distinct from the power-law response discussed above. In this case, temporal scale-invariance manifests as diverse, multiscale fluctuations distributed according to power laws. This type of scale-invariance affords a second route for a bifurcation to meet our definition of criticality.

To understand the origins of such noisy scale invariant dynamics, first consider the case when the linear terms of *f* (*x, µ*) dominate, i.e., when the activity is below the crossover *x*≪ *ξ*_*lin*_. Then, Eq. 1 becomes an Ornstein-Uhlenbeck (OU) process

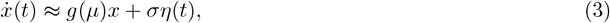

with a time scale that diverges as *µ* approaches *µ*_*b*_. This is another manifestation of critical slowing down. An OU process with infinite time scale is a continuous-time Brownian random walk, which is known to be temporally scale-invariant — a fixed point of temporal renormalization group theory^28^— with powerlaw power spectrum^37,38^, and power-law distributed first passage times^39–41^. Thus, unlike the noise-free deterministic case, the purely linear system with noise becomes scale invariant and conforms to our definition of criticality.

However, Brownian dynamics will inevitably wander beyond the crossover *ξ*_*lin*_; the greater the noise amplitude *σ*, the faster they will wander. Eventually, this results in *x* ≫ *ξ*_*lin*_ and Eq. 1 becomes

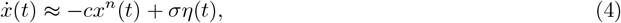

This equation becomes exact at the bifurcation, where the linear term is zero. Here, the nonlinear term −*cx*^*n*^(*t*) pushes *x* back towards *x*\*, keeping the Brownian dynamics from wandering too far. Likewise, if a stimulus kicks the activity well above the crossover *x* ≫ *ξ*_*lin*_, the nonlinear terms of in *f* (*x, µ*_*b*_) dominate, as in the deterministic case, and the power-law relaxation can become apparent again, albeit with noise added. Thus, for noisy systems, dynamics are governed by a competition between the nonlinear drift towards *x*\* and noise causing diffusion away from *x*\*. This competition entails another crossover scale *ξ*_*noise*_ that separates drift-dominated and diffusion-dominated regimes (see Fig. 3B); *ξ*_*noise*_ is revealed by equating their characteristic times (see Appendix A)

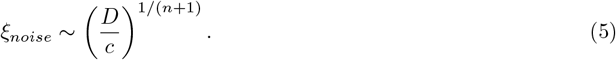

In other words, the relative size of the noise amplitude *σ* and the nonlinear term *cx*^*n*^(*t*) determine whether the dynamics exhibit power-law decrease towards *x*\* or power-law distributed Brownian fluctuations.

Thus, we arrive at a central finding. In general, there are two sources of scale-invariance that can give rise to critical dynamics in dynamical systems with noise: 1) non-linear, deterministic power-law response to stimulus and 2) linear, stochastic Brownian motion-type fluctuations. In the linear systems, only the second source is relevant; Brownian fluctuations are unbounded, producing true random walks at the boundary between stability and instability. In nonlinear systems, however, the two types of scale-invariance coexist with the nonlinear power-law drift keeping the Brownian fluctuations from wandering too far.

### Empirical signatures and predictions for experiments

The most common empirical approaches to test for criticality in neuroscience assess scale-invariance in the fluctuations of ongoing neural activity, similar to the Brownian fluctuations that emerge near bifurcations in low-dimensional systems with noise. Methods include neuronal avalanche analyses^25,42–44^, crackling noise scaling laws^24,45,46^, detrended fluctuation analyses^47,48^, power-spectra^49^, temporal renormalization group approaches^28^, susceptibility^50^, and more^3^. This raises the question of whether the underlying bifurcation can be identified by examination of avalanches that reflect Brownian criticality.

To address this, we performed avalanche analyses on time series generated by simulating Eq. 1 (see Appendix D). Briefly, an “avalanche” begins when activity rises above a threshold and ceases when activity falls below (Fig. 5A). For a long time series, thousands of avalanches are obtained, and probability distributions of avalanche size and duration are analyzed. In both highand low-dimensional systems, it is expected that these distributions have a wide range of power law scaling, thus indicating scale invariance^25,40^. Moreover, the exponents of the size and duration distributions should be related according to the crackling noise scaling law^46,51^. We performed simulations and avalanche analyses for three bifurcations: first, a linear system (purely Brownian with no drift), second, the supercritical pitchfork bifurcation, and third, the transcritical bifurcation (Fig. 5).

**Figure 5:**
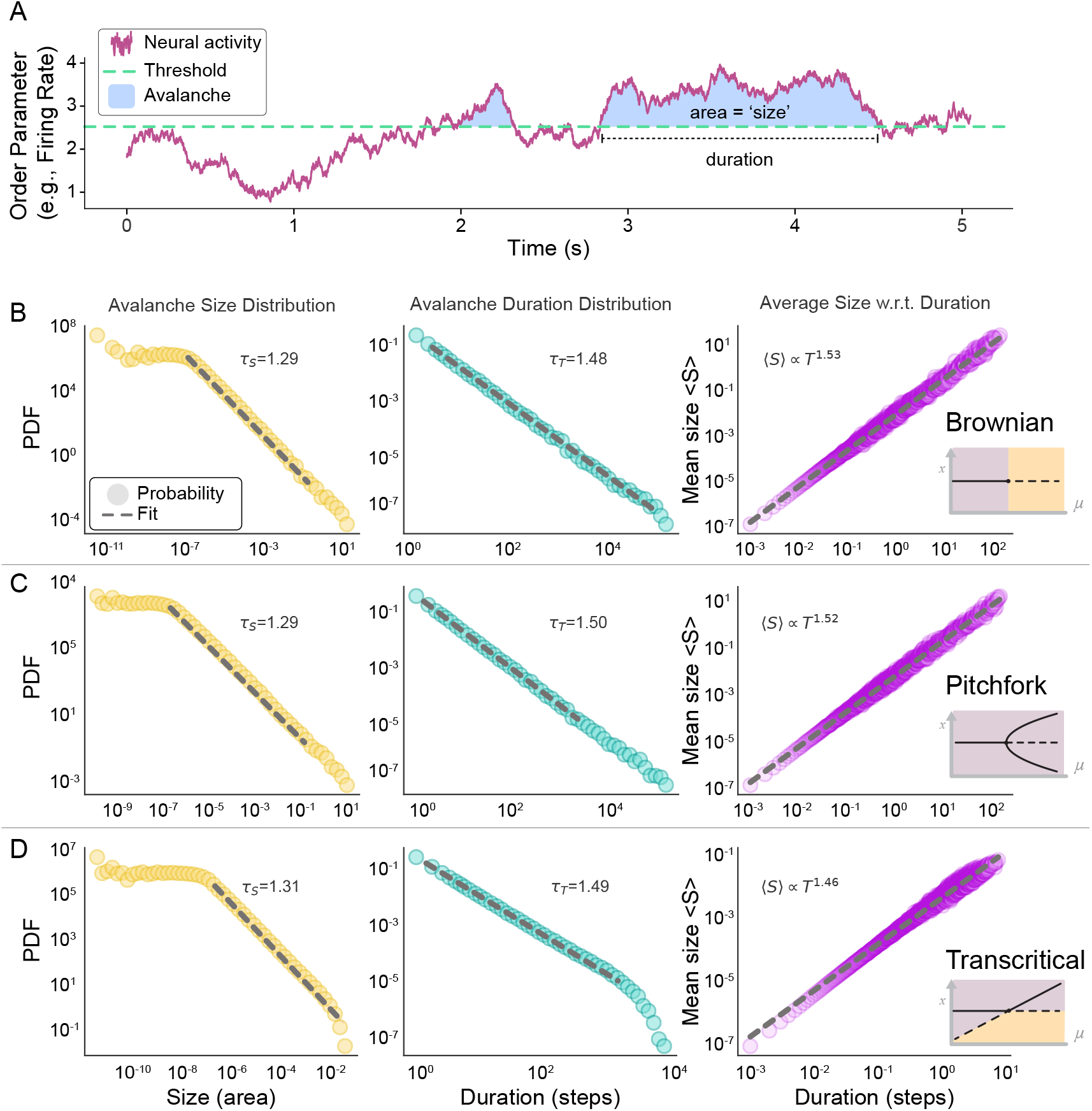
At a bifurcation, noise-driven fluctuations produce scale-invariant avalanches. **(A)** Neuronal avalanches are fluctuations in neural activity (purple) that cross above a threshold (dashed green). Avalanches are typically quantified in terms of their size (blue shaded area) and their duration. **(B)** In the *pure* Brownian case, i.e., a linear dynamical system at its bifurcation, there is no tendency of the system to relax back to a fixed point; spurred by noise or perturbation, dynamics can wander with no bounds. Brownian fluctuations produce avalanches with power-law distributed sizes (left) and durations (middle) and conform to the crackling noise scaling law (right). Together, these three aspects of temporal scale invariance are commonly used to empirically support the criticality hypothesis in neural systems^24,25,50,52^. **(C)** Tuned to the bifurcation in a supercritical pitchfork, the only difference is the continual drift back toward the fixed point — the system exhibits a central tendency and cannot wander without bound. Here, the addition of additive Gaussian noise results in avalanches following power laws nearly identical to the linear case. **(D)** Same as B and C, but in a transcritical model (with *x >* 0 enforced). Each distribution represents the concatenation of 10 random runs. Dashed gray line indicates the region of the distribution fit by a power law, whose exponents are displayed in each panel.

In the absence of large amplitude perturbations on top of the noise, we expected our simulations of all three systems to generate approximately Brownian dynamics. Accordingly, size and duration distributions exhibited many decades of power-law scaling and the exponents for the three systems were nearly identical (Fig. 5) and close to known exponents for Brownian motion[40]. Further, each satisfied the crackling noise scaling law (Fig. 5, right column). These results suggest that experimentally observed avalanches in awake animals^42,52^, which exhibit behavior similar to our simulations, might be effectively modeled by low-dimensional bifurcations. However, the nonlinear structure of the bifurcation is effectively invisible to standard avalanche analyses.

Given the dominance of Brownian criticality in avalanche distributions, it is important to ask how the bifurcations underlying neural data can be discerned in the face of inevitable noise. Neither avalanches nor other standard approaches assess power-law relaxation, which is an independent source of criticality and varies according to the bifurcation type. As explained above, the response to a brief, strong stimulus sufficiently above the crossover scale (*x*≫*ξ*_*noise*_, Fig. 3B and Eq. 5) should expose both the underlying bifurcation and how close *µ* is to the bifurcation. Specifically, at a nonlinear bifurcation, the relaxation exponent should indicate the bifurcation type (see Appendix A). To test this, we ran numerical simulations of the supercritical pitchfork with additive Gaussian noise subject to a series of stimuli below, at, and above the bifurcation (*µ* = −1, 0, and 1; Fig. 6).

**Figure 6:**
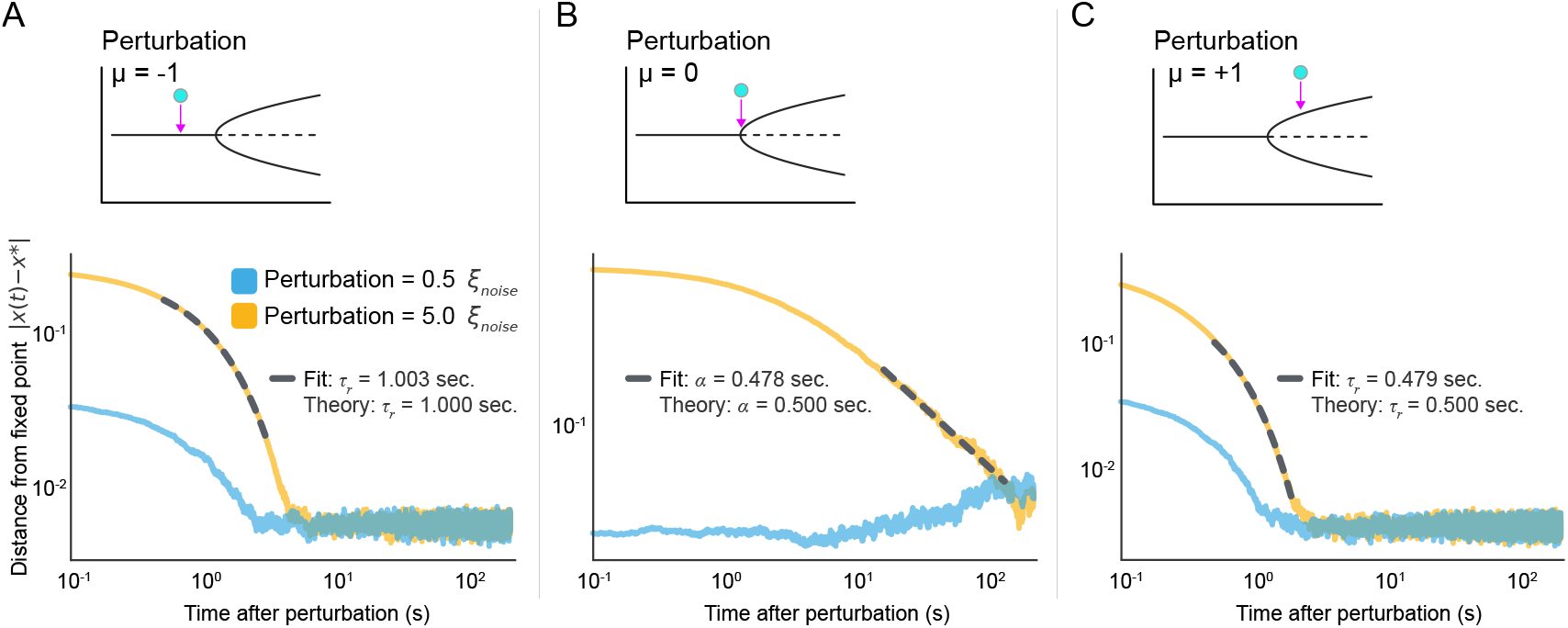
Strong perturbations reveal proximity to and type of bifurcation. The Brownian dynamics for *x ≪ ξ*_*noise*_ are largely independent of the type of bifurcation. Here we ask whether response to a strong perturbation can be more revealing. We perturbed a supercritical pitchfork below, at, and above the bifurcation in two contexts: perturbations *less than* the noise crossover scale (0.5*ξ*_*noise*_) and *greater than* the scale (5*ξ*_*noise*_). **(A)** Below the bifurcation (*µ < µ*_*b*_), response to perturbations *below* the noise crossover scale (blue) or *above* the crossover scale (gold) both exhibit exponential decay. The exponential fit of the gold curve is shown in dashed gray, and compared to the theoretical value. **(B)** Same as **A** but for perturbations at the bifurcation (*µ* = *µ*_*b*_). Dashed gray indicates power law fit (not to be confused with *τ* of **A** and **C**). **(C)** Same as **A** but for perturbations above the bifurcation (*µ > µ*_*b*_). Each trace represents an average over 200 repeated simulations.

As predicted, the system response to a weak stimulus (0.5*ξ*_*noise*_) did not reveal the nonlinear structure of the bifurcation. Stronger stimulation (5*ξ*_*noise*_) at *µ* = −1 resulted in a response that decayed exponentially with relaxation time *τ*_*r*_ ≈ 1 seconds. Similarly, perturbing at *µ* = 1 resulted in another exponential relaxation with relaxation time *τ*_*r*_ ≈ 1*/*2. Finally, tuned precisely to the bifurcation, the perturbation decayed as a power law with exponent *α* = 1*/*2. Each of these is consistent with the theoretical description of the supercritical pitchfork (Fig. 6 and Appendix A).

Taken together, these results demonstrate that, even in the presence of Brownian criticality at small scales, the underlying bifurcation *can* be extracted by examining the response to a sufficiently intense stimulus. This directly informs mathematically rigorous experimental design in neuroscience via the application of pulsed perturbations. Such an approach would offer a powerful method of empirically reconstructing the bifurcation(s) underlying neural circuits, thus directly reconciling theory and biology. Crucially, stimuli must drive activity substantially beyond the amplitude of ongoing fluctuations (beyond *ξ*_*noise*_) and be separated by enough time to observe complete relaxation (Fig. 6B).

### Which critical bifurcations are biologically plausible explanations of the brain’s computational regime?

Thus far, we have only explored perturbation and noise in the order parameter *x*, implicitly assuming that the control parameter *µ* is constant. In real world systems, noise impacts not only the order parameter (e.g., neuronal activity), but also control parameters that tune a system’s state. Control parameters are not fixed; neuromodulation, synaptic plasticity, and other factors exhibit noise and continuously influence the brain’s operating conditions. Thus, if a critical bifurcation accounts for the information processing capacity of a biological network, slight variation in *µ* must not lead to catastrophic failure, i.e., activity blowing up to ±∞ or the system jumping to a far away state. Robustness to variability in the control parameter is not guaranteed. In Fig. 4, any variation in *µ* or *x* that places the system in the unstable (yellow) region of the bifurcation diagram can cause catastrophic failure. This occurs for several bifurcation types: saddle node, unbounded transcritical, subcritical pitchfork, subcritical Hopf, and linear.

In some cases, instability is directional; in saddle nodes, perturbations in *x* restricted to the stable side will not suffer catastrophic failures. From the other direction, perturbations cause the system to fly away, either jumping to a distant, separate fixed point, or to infinity (e.g., subcritical pitchfork with saddles in Fig. 2B or isolated saddle node in Fig. 4 respectively). Finally, linear models are infeasible, because, at the bifurcation, they have no tendency to return following any change in *x*. Even the unbiased random walk can drift to infinity. Although not explosive, this is nonetheless not realistic if *x* represents any aspect of neural activity. Thus, the list of bifurcations that both generate criticality and are biologically robust includes: supercritical pitchfork, supercritical Hopf, and transcritical with *x >* 0. These are unified by continuity along stable fixed points, which bears stark similarity to the requirement of a continuous phase transitions for criticality in high dimensional systems (*N*→ ∞). Fig.4 summarizes these considerations. We note that some aspects of neuronal phenomena benefit from invoking instability, such as spike generation and sleep/wake transitions. However, in models of sleep/wake transitions (e.g.), the blow-up associated with a saddle node is mitigated by including another stable fixed point that the system can fall into (i.e., sleep or wake – the other state)^36,53^.

### Information and Bifurcations

Broadly considered, there is selective pressure for almost all neural systems to achieve a regime that maximizes general information processing. Criticality has been suggested as general principle of optimal computation, particular for flexible systems that adapt to a complex and *a priori* unpredictable information^3–5^. However, research about optimal information processing near criticality has been conducted almost exclusively in high-dimensional systems (*N* ≫ 1)^2,3,54–56^. This leaves open the question of whether a critical bifurcation represents an analogous informational optimum in low dimensional systems.

As argued above, noiseless deterministic systems are not biologically feasible. Thus, we sought to understand how noisy, low dimensional dynamical systems process information. To address this, we measured mutual information between a brief stimulus and the resultant response over time. Specifically, if a random variable *X* is given a brief pulse perturbation *δX* at a given time such that *X* → *X* + *δX*, the mutual information between the stimulus and state of the system at a later time (*τ*) can be written as

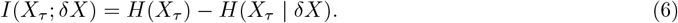

where 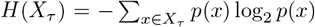 is the Shannon entropy of *X*_*τ*_, and *H*(*X*_*τ*_ | *δX*) is the conditional entropy of responses *X*_*τ*_ dependent on the perturbation *δX*^57^. We estimated both *H*(*X*_*τ*_) and *H*(*X*_*τ*_ *δX*) from the ensemble distributions obtained across stochastic realizations of the system under identical conditions, thus quantifying how much knowledge of the stimulus reduces uncertainty about the future state (see Appendix D). For each value of the bifurcation parameter, we let the system settle for *T*_settle_ = 100 *s* before applying the pulse perturbation. Then, *n* = 500 trials were run with different perturbations *δX* = {0.05, 0.1, 0.2, 0.3, 0.5, 1} and evolved for delays *τ* ∈ [1, 19]*s*. Analyses were conducted with *D* = 5 × 10^−5^, thus all perturbations excluding *δx* = 0.05 were above the crossover scale *ξ*_*noise*_ at the bifurcation. We tested supercritical pitchfork, transcritical, and saddle-node bifurcations as well as the linear model. To avoid metastability artifacts for the pitchfork, we used a within-basin constraint to study mutual information locally near a fixed point. Similarly, for saddle-node and transcritical bifurcations, we enforced *x >* 0 to avoid runaway activity. In all four instances, mutual information grows as the bifurcation is approached (Fig. 7). It is noteworthy that the rate of growth is symmetrical in the transcritical model, but not the pitchfork (Fig. 7A,B), reflecting the underlying geometry of the bifurcation. In contrast with purely Brownian systems, the other three bifurcations (which exhibit a consistent drift towards the fixed point) exhibit nonlinear decay in mutual information over time. The Brownian system, however, maintains stimulus information for extended periods (Fig. 8). Taken together, these results are consistent with previously observed information maximization near criticality in large *N* systems^54,56,58^.

**Figure 7:**
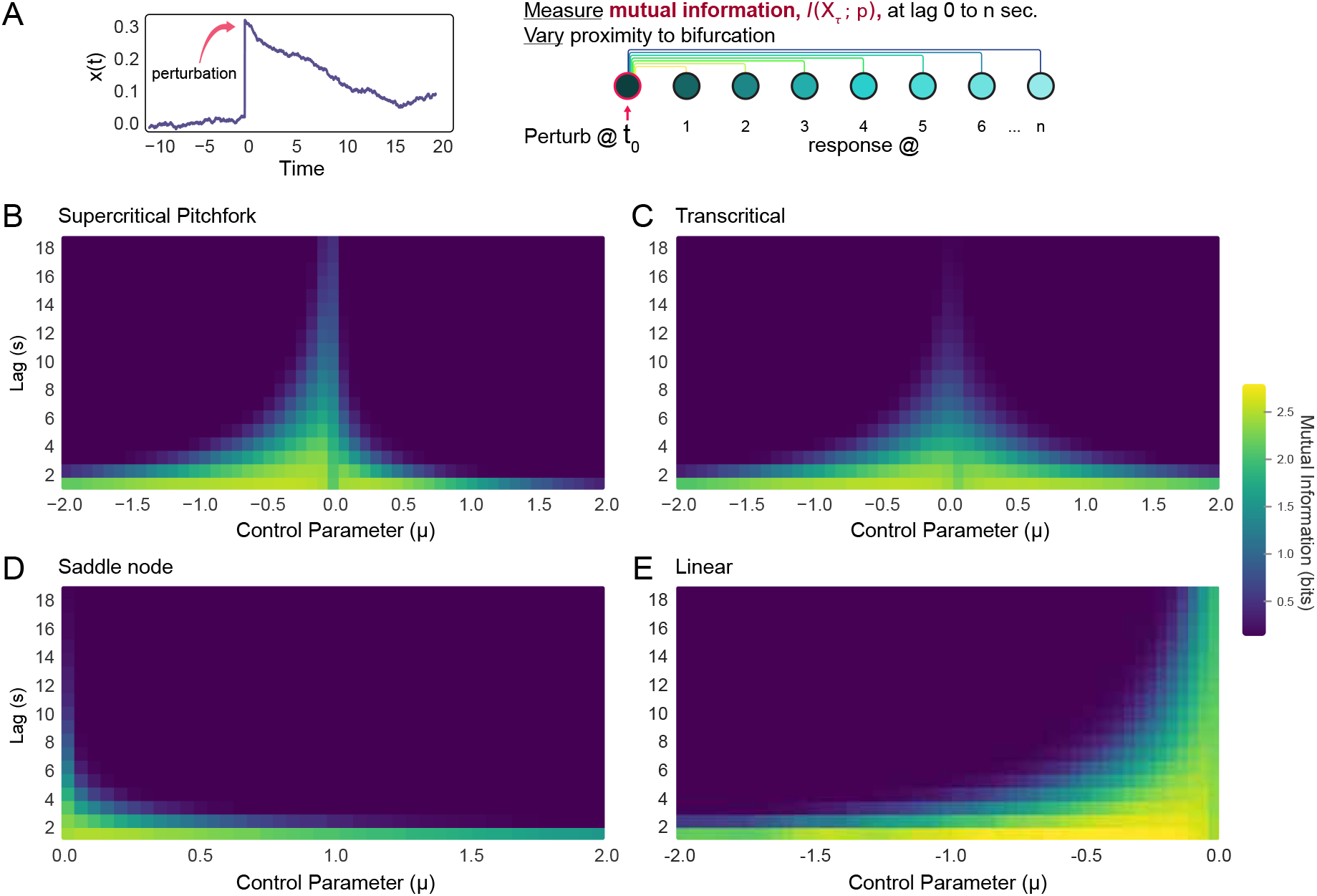
Temporal mutual information is maximized at critical bifurcations. **(A)** Graphic illustration of numerical analysis. (Left) Sample data showing the order parameter of a system before, during, and after a perturbation at time = 0. (Right) Following perturbation, mutual information — *I*(*X*_*τ*_; *δX*)— was calculated for 20 different lags into the future. In other words, how long did information about the perturbation persist in the system? We performed this numerical analyses in **(B)** the supercritical pitchfork, **(C)** transcritical bifurcation, **(D)** saddle-node bifurcation, and **(E)** a linear OU model. For B-E, 500 trials comprising a range of perturbations were conducted.

**Figure 8:**
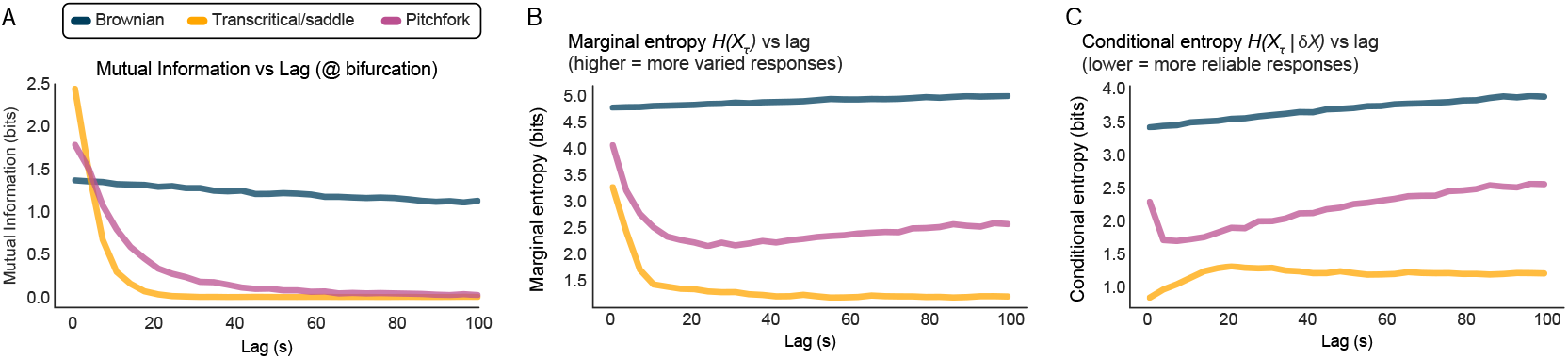
Mutual information, marginal, and conditional entropies at the bifurcation. **(A)** Mutual information versus lag *at the bifurcation* for purely Brownian criticality, the transcritical/saddle (at bifurcation, these exhibit the same polynomial, see Eq. 13), and pitchfork..**(B)** Marginal entropy (the diversity of responses) across a range of lags in Brownian, transcritical/saddle, and pitchfork systems set to the bifurcation (*µ* = 0). Higher values indicate more varied responses to the same input. **(B)** Same as **A** but conditional entropy (the degree of randomness given knowledge of the perturbation). The reliability of responses decreases as conditional entropy increases.

There is ongoing debate about whether linear/Brownian criticality is the source of scale invariance in the brain^31,59^. Linear systems lack stability and reliability as their time-constants approach very large values; neural models employing only Brownian motion are free to reach infinite firing rates and variance, for example^60^. Consistent with this, we find that responses in purely Brownian systems are not reliable as they never start from the same initial conditions. This is reflected by the large conditional entropy in Brownian criticality, indicating substantial variability in responses to the same inputs (Fig. 8B). Summarily, while purely Brownian criticality maintains information with high fidelity across extended delays, our results suggest that it is not likely to single-handedly account for scale invariance in biological systems.

### Do previously studied neural models exhibit critical bifurcations?

Criticality has been proposed as a unifying principle of brain function^3^. If true, models that have accurately captured significant aspects of brain function should be constrained by the same fact. However, given the preponderance of low-dimensional dynamical models in neuroscience and the lack of a definition of criticality in this context, it has been impossible to address this. We next divided historically significant models into six categories and asked if the underlying bifurcations are critical.

#### Single-neuron models

The dynamics of spiking neurons, particularly transitions between resting, tonic activity, and bursting, are governed by low-dimensional systems that exhibit characteristic bifurcations. Here, the order parameter is membrane potential, and control parameters are typically intrinsic or synaptic currents. Prominent models include those of Hodgkin and Huxley^13^, FitzHugh and Nagumo^61,62^, and Morris and Lecar^63^. Eugene Izhikevich^64^ organized single-neuron excitability models into four canonical types defined by the bifurcation responsible for the onset of repetitive firing: 1) integrator and bistable, 2) resonator and bistable, 3) integrator and monostable, and 4) resonator and monostable. These employ saddle-nodes, subcritical Hopf, saddle node on invariant circle, and supercritical Hopf bifurcations respectively. Our results show that the saddle node and supercritical Hopf are critical, whereas the subcritical Hopf is not.

Another historically prominent description of single neurons is the firing rate model^9,65–67^, which may be expressed as

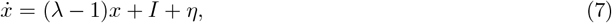

where *λ* is a control parameter representing effective gain or excitability, *I* is external input from neighbors or environment, and *η* is Gaussian noise. For *λ <* 1 the system is linearly stable, while for *λ >* 1 it becomes unstable, with the relaxation time diverging at the critical value *λ*_*c*_ = 1. Because dynamics are linear in the order parameter, no nonlinear restoring term is present, and thus the system does not display power-law relaxation characteristic of deterministic criticality. Instead, near the critical point, dynamics reduce to pure Brownian criticality in which scale-invariant fluctuations dominate.

Taken together, these results demonstrate that the activity of a single neuron can be modeled *without* invoking a critical bifurcation. In contrast to the single neuron, networks involve multiple, connected nodes and can be as small as two and as large as 10^11^ neurons. The remaining four categories comprise models of collective interactions across varying scales.

#### Discrete-state network models

Discrete-state network models comprise connected neurons that can occupy binary states (0/1 or ± 1). These evolve according to deterministic or stochastic update rules based on weighted inputs from neighboring units. A famous example is the McCulloch and Pitts model^68^, which was foundational for early neural computation and inspired architectures such as the perceptron^69^— the first artificial neural network capable of learning from data. In neurobiology, such models have been effective in studying avalanche dynamics and power laws^24,70^.

Depending on their construction, both deterministic and probabilistic discrete-state network models can display a variety of underlying bifurcations in collective activity, including pitchfork bifurcations^71^, transcritical bifurcations^18,24,25,50^, saddle nodes^71^, and supercritical Hopf bifurcations^72^. Feedforward architectures without recurrence, such as finite perceptrons, do not exhibit local bifurcations; however, in the limit of infinitely many layers or probabilistic propagation across layers, they can reach a critical point separating quiescent and self-sustaining activity, displaying a transcritical bifurcation^12,17,18,25^. In simple terms, recurrent models contain a critical bifurcation, and feed-forward models approximate criticality in a large *N* system (where *N* is the number of layers).

#### Continuous-state network models

The most direct application of dynamical systems theory to neuroscience is the representation of neural activity by ordinary differential equations. These can be generalized as

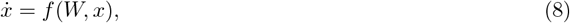

where *x* ∈ ℝ^*n*^, *W* is the connectivity matrix, and *f* (·) is a continuous function (noise can be included). Two main models emerge: rate-based and phase oscillator models.

Rate models include the influential Wilson-Cowan mean-field model^15^ as well as many recurrent neural networks frequently used to model continuous-state neural data from meso-scale recording modalities^73^. If weights are exclusively positive, these models map to the mean-field Ising model with its respective critical phenomena^74^, i.e., the supercritical pitchfork bifurcation^30^. Allowing for inhibitory interactions, other bifurcations, including saddle nodes and supercritical Hopf bifurcations, may appear^15^. Furthermore, artificial deep neural networks, which are often rate models, exhibit transcritical bifurcations^75^. This analysis reveals that continuous-state rate models all involve critical bifurcations.

Continuous-state networks can, instead of rate, also be described as coupled oscillators. In this case, one arrives at the the Kuramoto model^21^ which captures synchronization phenomena and phase transitions. The Kuramoto model has been used to investigate neural synchronization and rhythm generation and displays supercritical Hopf bifurcations^76^. This is also a critical bifurcation.

Another class of continuous state model comprises the spiking neural network, which is a hybrid of discrete and continuous states. Spiking neural networks track membrane potential (continuous state) and use thresholds to reset after an action potential (discrete state). In some respects, these models involve building a network out of the single neuron models discussed above. The most common spiking model is the leaky integrate-and-fire (LIF) neuron which dates back to Louis Lapique in the early 1900’s^20^. Networks of LIF neurons display a wide range of bifurcations including supercritical pitchfork, transcritical, saddle node, and supercritical Hopf^17,26,77^. A thorough mathematical analysis of the phase space of LIF networks is still lacking, but it is well known that these networks exhibit many of the same signatures of criticality as discrete-state models, giving rise to avalanche power laws.

A major class of models used for meso and macroscale phenomena are *field models* which describe neural activity as continuous spatiotemporal fields and allow thorough analysis of correlation functions. Classical models include the Wilson-Cowan^78,79^ with spatial structure, and the Amari model^80,81^. Mean field models capture pattern formation, traveling waves, and critical phenomena amenable to field-theoretic methods (*N*→ ∞) and renormalization group analysis^82^. These models map directly to the transcritical bifurcation as well as other more diverse but still critical bifurcations^83^. Once again, each of these is a critical bifurcation.

Collectively, these analyses reveal that, beyond the level of the single neuron, all examined models of neural interactions invoke dynamical systems that contain critical bifurcations (Table 1). This adds further support to the hypothesis that criticality, or at least the capacity to be tuned to a critical point, is a unifying computational requirement of any model of the brain, assuming only that a brain is larger than one neuron.

**Table 1:** Historically relevant neural models and their characteristic bifurcations. While this table is not exhaustive, many specialized models employed throughout the literature and not specifically considered here are variants of the six forms presented in this table. In the list of typical bifurcation (right column), those that are critical are italicized for ease of classification.

| Model Type | Examples | Typical Bifurcations |
| --- | --- | --- |
| **Single-neuron models** |  |  |
| Deterministic spiking | Izhikevich; Hodgkin–Huxley; Morris–Lecar; FitzHugh–Nagumo | *SNIC*; *saddle-node*; *supercritical* or subcritical Hopf |
| Linear rate (driven) | Leaky integrator; linearized firing-rate model | Change in stability at $\lambda_c = 1$; *Brownian criticality* |
| **Population and network models** |  |  |
| Discrete-state | McCulloch–Pitts; probabilistic automata; branching models | *Transcritical*; *pitchfork*; *supercritical Hopf* |
| Firing-rate | Wilson–Cowan; recurrent neural networks (RNNs) | *supercritical Hopf*; *supercritical pitchfork*; *saddle-node* |
| Phase-oscillator | Kuramoto; Winfree | *supercritical Hopf* |
| Neural field | Amari; spatial Wilson–Cowan; stochastic neural fields | *supercritical Hopf*; *Turing–Hopf*; *saddle-node*; *transcritical* |

## Discussion

A proposed unifying principle of neural function must be reconcilable, at the very least, in both large and small scale models of the brain, as each approach can yield significant explanatory insight. More specifically, the brain’s computational power has been suggested to arise from operating near critical points, yet many of our most successful mechanistic models are low-dimensional dynamical systems where thermodynamic phase transitions — and thus criticality — have not been well-defined. In addition to fueling confusion when comparing different approaches to modeling the brain, this disconnect has prevented an “apples-to-apples” mechanistic comparison of models across scales. We set out to resolve this by developing a rigorous definition of criticality for dynamical systems that decouples scale-invariance from the spatial domain. Mathematically, the same features of scale invariance that emerge in spatially distributed systems at criticality can be recapitulated in temporal properties of low-dimensional systems. Applying this framework, we demonstrate that canonical bifurcations in neuroscience — transcritical, supercritical pitchfork, saddle-node, and supercritical Hopf — exhibit true criticality characterized by power-law relaxation and Brownian fluctuations. We further show that information processing capacity, quantified by mutual information, is highest precisely at these critical bifurcations. Consistent with this, we find that all historically successful models of neural communication, but not necessarily models of spike generation, operate at or at least contain critical bifurcations. These results demonstrate that criticality is not an emergent property exclusive to many-body systems, but is a fundamental organizing principle accessible to systems of any size. This is attractive, as it appears to neatly harmonize decades of idiomatic modeling approaches under a single computational framework.

In nature, biology is constrained by two facts. First, noise (both internal and external) is inevitable, and second, infinity is non-viable. Within these constraints, and because neural systems operate far from equilibrium — that is, they continually dissipate energy to maintain functional organization — informationally optimal brain activity must be defined by two intertwined regimes. The first, Brownian criticality, emerges when stochastic fluctuations act on a system poised near a bifurcation, producing scale-free fluctuations that reflect a random-walk through the local landscape. This regime is remarkable for its preservation of information over extended delays but lacks intrinsic restoration — each perturbation shifts the baseline without a counteracting force. As a result, purely Brownian dynamics are doomed to wander away and lack reliable input/output relationships. Each of these features is incompatible with metabolic and informational stability, which, in turn, are first-order requirements for a viable brain. Our results indicate that the nervous system may pair random-walk exploration with an independent, complementary source of scale invariance that provides a dissipative (i.e., homeostatic) structure. This deterministic criticality imposes a nonlinear restoring flow that bounds fluctuations and is itself another informational optimum. This principal finding leads to the conclusion that there are two sources of scale invariance in any biologically plausible network. In other words, the brain maintains itself near a critical manifold where power-law dissipation bounds the fluctuations driven by inevitable noise, maintaining high information capacity without violating energetic or stability constraints.

It is reasonable to suggest that there is a strong selective pressure on brains to maximize computational capacity^3^. Our results reveal a striking convergence: information-theoretic measures are greatest at criticality regardless of whether one approaches from the perspective of many-body phase transitions or low-dimensional dynamical systems. The mutual information *I*(*X*_*τ*_; *δX*) between initial perturbations and future states peaks precisely at the critical point for the pitchfork, transcritical, and saddle-node bifurcations, mirroring the enhanced information transmission observed in extended systems near thermodynamic criticality^54–56,84^. This convergence is not coincidental but reflects a deeper principle: criticality, whether manifest as a phase transition in infinite-dimensional systems or as a bifurcation in low-dimensional dynamics, creates the conditions necessary for complex computation. At this point, a brain becomes maximally sensitive to inputs while avoiding instability, achieving what has long been suggested as a computational predicate for optimal function^1,3,24,25,85–87^. Crucially, this demonstrates that the computational benefits attributed to criticality — enhanced dynamic range, optimal information storage and transmission, etc. — are not artifacts of any particular modeling framework but fundamental properties of critical points themselves, accessible through multiple mathematical formalisms all pointing to the same underlying computational principle.

Remarkably, nearly every canonical model in theoretical neuroscience—from Wilson-Cowan population dynamics to Kuramoto oscillators, from synaptic plasticity rules to slow-wave sleep models—contains bifurcations that meet our definition of criticality. This is striking: these models were developed independently over decades to explain disparate phenomena, yet they converge on the same mathematical structures that give rise to criticality. The notable exception is instructive: models of spike generation, such as the Hodgkin-Huxley equations and integrate-and-fire neurons, typically operate through saddle-node bifurcations on invariant circles or homoclinic bifurcations, some of which our analyses show do not support criticality. This distinction makes biological sense — individual spike generation requires reliable, stereotyped responses rather than scale-free flexibility. However, once we move beyond single spikes to consider population activity, oscillations, plasticity, or any form of collective computation, critical bifurcations appear to become the rule rather than the exception. This dichotomy suggests that while neurons must maintain robust spiking mechanisms for reliable signal transmission, the computational flexibility required for learning, adaptation, and information processing is achieved from dynamics related to scale invariance.

This framework makes specific, falsifiable predictions that distinguish critical bifurcations from other dynamical regimes and from generic claims about scale-invariance. Most directly, neural systems operating near critical bifurcations should exhibit power-law relaxation following perturbations, with the relaxation following *δx* ~ *t*^*−α*^ where the exponent *α* depends on the bifurcation type — a prediction that, surprisingly, has never been directly tested in neural systems. By delivering controlled perturbations of varying magnitude and measuring the full relaxation trajectory back to baseline, it should be possible to measure the presence (or absence) of deterministic/emergent criticality. Systems at a critical bifurcation should reveal relaxation times that follow *δx*(*t*) ~ *t*^*−α*^, with *α* = 1 for transcritical bifurcations and *α* = 1*/*2 for pitchfork bifurcations.

Furthermore, our analysis makes specific predictions for avalanche distributions. If fluctuations in neuronal activity are purely Brownian, then the exponents will be *τ*_*T*_ = 3*/*2 for duration, and *τ*_*S*_ = 4*/*3 for size. On the other hand, if the underlying fluctuations emerge from collective interactions, then the exponents will belong to a class of models treated in the literature of phase transitions^30,88^. For transcritical bifurcations relevant to neuronal avalanches and mean-field interactions, avalanche exponents should be *τ*_*S*_ = 3*/*2 and *τ*_*T*_ = 2, corresponding to the mean-field directed percolation universality class. These precise exponent values, combined with the requirement for diverging relaxation times and the specific relationship between scaling exponents, provide strong constraints that can definitively test whether a neural system operates at a critical bifurcation or merely exhibits a heavy-tailed distribution arising from other mechanisms. Modern experimental techniques combining optogenetics for precise perturbations with high-density recordings make such tests immediately feasible, offering a clear path to validate or refute the criticality hypothesis in specific neural circuits and contexts.

For the sake of conciseness, we left many features of critical phenomena aside. Of particular relevance to this work is the theory of universality classes which are defined by the set of critical exponents of a given model^89^. Many of these critical exponents are only observed in the neighborhood of the critical point, i.e., in the limit of *µ*→ *µ*_*b*_. Extending the present framework to classify the full extent of low-dimensional normal forms into universality classes (or families in the absence of spatial degrees of freedom) would further unify bifurcation theory and thermodynamic phase transition theory. Another line of development includes the problem of adaptability and feedback loops in coupled dynamical systems, which can give rise to selforganized criticality, a fundamental problem in biology. Despite a vast literature addressing particular cases, a systematic general theory remains largely unsolved^90^. Expanding this framework and its relationship to information processing could move us towards a foundational theory of brain dynamics, and, more broadly, of biological organization.

## Acknowledgments

The authors thank Cyprien Tamekue and Gemechu Bekele Tolossa for their incisive comments on bifurcation theory.

## Appendix

### Regime crossovers and response to perturbations

We consider the dynamics

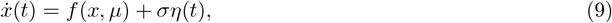

which covers all the dynamical systems discussed in the main text. Setting *σ* to zero recovers all the deterministic cases. To study the effect of a perturbation, we approximate the dynamical system using a Taylor expansion of *f* (*x, µ*) nearby a fixed point *x* = *x*\*+ *δx*, where *δx* is the perturbation^91^.

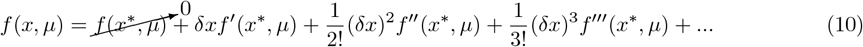

where the primes indicate partial derivatives with respect to *x*, e.g., 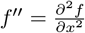. Since *x*\* is a fixed point, the first term *f* (*x*\*, *µ*) = 0 by definition. As discussed in the main text, the system dynamics can be understood by considering two different crossovers, *ξ*_*lin*_ and *ξ*_*noise*_. These crossovers are defined by considering the relative importance of three quantities: 1) the noise amplitude *σ*, 2) the linear term in the Taylor expansion *δxf*′(*x*\*, *µ*), and 3) the nonlinear terms in the Taylor expansion.

The crossover *ξ*_*lin*_ is defined by the relative size of the linear term compared to the nonlinear terms. For example, consider the supercritical pitchfork, *f* (*x, µ*) = *µx* − *x*^3^. Then the Taylor expansion near the fixed point 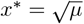 is

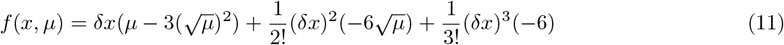

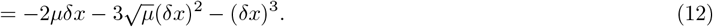

From this, we can derive the condition for *δx* resulting in a linear term that is much larger than the quadratic term, 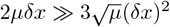, which implies 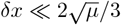 In other words, the crossover 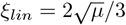. Similarly, another crossover could be derived to determine when the third order term dominates the second order term, but it is beyond the scope of the present work. Table 2 summarizes the crossover values of *ξ*_*lin*_ for the normal form bifurcations we considered in the main text.

**Table 2.** Summary of crossover values *ξ*_*lin*_. For *δx ≪ ξ*_*lin*_, linear dynamics dominate. For *δx ≫ ξ*_*lin*_, nonlinear dynamics dominate.

| Bifurcation type | $f(x, \mu)$ | Fixed points | $\xi_{lin}$ |
| --- | --- | --- | --- |
| linear | $\mu x$ | $x^* = 0$ | ∞ |
| transcritical | $\mu x - x^2$ | $x^* = \{0, \mu\}$ | $\mu$ |
| saddle node | $\mu - x^2$ | $x^* = \pm\sqrt{\mu}$ | $\pm 2\sqrt{\mu}$ |
| supercritical pitchfork, Hopf | $\mu x - x^3$ | $x^* = \{0, \pm\sqrt{\mu}\}$ | $\{\sqrt{\mu}, \pm 2\sqrt{\mu}/3\}$ |

Another important crossover, *ξ*_*noise*_, arises when considering the relative importance of Gaussian noise and the nonlinear terms in the Taylor expansion. The noise amplitude *σ* = 2*D* is the standard deviation of the noise where *D* is the diffusion constant. As discussed in the main text, near a bifurcation the linear terms become negligible and we’re left with

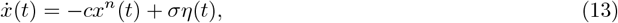

where *n* is the lowest nonvanishing polynomial and *c* is a constant. Then, we can find the crossover scale *ξ*_*noise*_ by first balancing the time scale of deterministic drift *τ*_drift_ and the time scale of noise driven diffusion *τ*_diff_. The time scale for a displacement of order |*x*| due to deterministic drift is

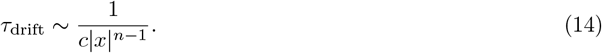

The time scale of diffusion, assuming Brownian motion, can be derived from the variance ⟨(*x*(*t*) − *x*(0))^2^⟩ = 2*Dt*. The time to diffuse a distance of order |*x*| is

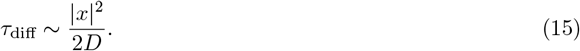

We define *ξ*_*noise*_ by identifying the condition on *x* that makes these two time scales equal

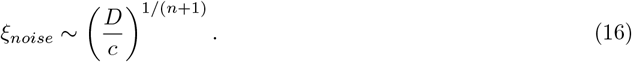

#### Linear regime: exponential relaxation

Whenever *x* ≪ *ξ*_*lin*_ the dynamics operate in a linear regime. Hence, to first order approximation, we have

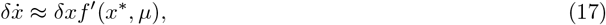

which has solution

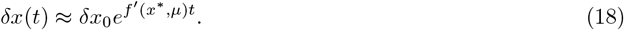

This indicates the temporal relaxation of a infinitesimal perturbation, thus the relaxation time is

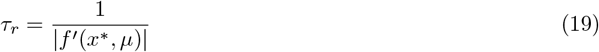

We can see that if *f*′(*x*\*, *µ*) *>* 0 the perturbation grows exponentially, whereas if *f*′(*x*\*, *µ*) *<* 0, the perturbation decays exponentially, whereas for *f*′(*x*\*, *µ*) → 0 we have *τ*→ ∞ which is known as *critical slowing down*. This occurs when approaching a bifurcation since in its normal-form, *f*′(*x*\*, *µ*_*b*_) = 0. In higher dimensional dynamical systems, this occurs when the real part of at least one eigenvalue goes to zero.

#### Nonlinear regime: scale invariant relaxation

Whenever *x* ≫ *ξ*_*lin*_ the dynamics operate in a nonlinear regime.

As the control parameter approaches *µ*_*b*_, the relaxation time diverges. At this point, the linear term vanishes (first derivative), and the leading non-linear terms becomes relevant. Hence, at the bifurcation, if the second order derivative is the leading nonlinear term, then

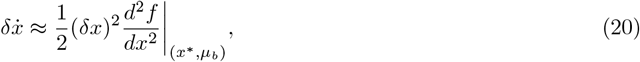

which, by separation of variables, leads to the solution^92^

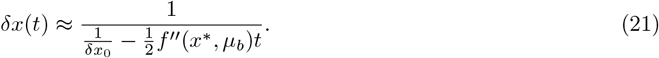

As *t* grows, we can see that it will overpower 1*/δx*_0_, so we have

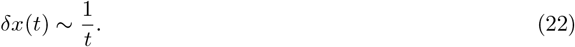

Note that from Eq. 21 we can see the perturbation has to be bounded depending on the sign of *f*^*′′*^(*x*\*, *µ*_*b*_) otherwise it blows up in finite time. This is why quadratic bifurcations such as the transcritical and saddle node need to be bounded.

From the derivations above we can see that the leading order derivative that remains beyond *f*′(*x*\*, *µ*_*b*_) will give the value of the exponent *α* described previously. In general, the solution can be represented as,

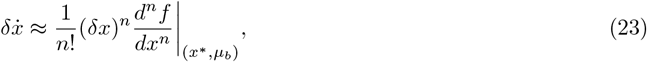

where *n* represents the leading order that remains due to the symmetries of the model and other considerations. Hence, we can find the value of the exponent *α* in terms of *n*. It is very simple to show this leads to the relationship

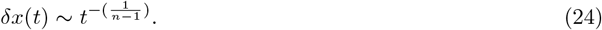

Hence, we can see that *α* = 1*/*(*n* − 1) for *n ≥* 2. As stated with the quadratic case, it is important to note that depending on the sign and symmetry of the *n*th derivative, the solution to the perturbation may blow up in finite time.

### Numerical simulations and analysis

Numerical simulations for dynamical systems under noise were carried out using the Euler-Mayurama method. The parameters were: time step Δ*t* = 0.001, noise standard deviation *σ* = 0.01. For the avalanche distribution calculations we used the median as a threshold and collected avalanches from multiple runs with same parameters (10 runs).

For the information theory section, the parameters, time step, and noise STD were the same as previously. For each value of the bifurcation parameter, we let the system settle for *T*_settle_ = 100 *s* before applying the pulse perturbation. Then, *n* = 500 trials were run with different perturbations *p* = [0.05, 0.1, 0.2, 0.3, 0.5, 1] and evolved for delays *τ* ∈ [1, 19]*s*. To estimate *I*(*X*_*τ*_; *p*), we used a histogram method with *n*_*bins*_ = 60 equally spaced bins spanning the pooled range [*x*_min_, *x*_max_] of all post-kick trajectories for each (*µ, τ*).

